# A cost-effective and scalable approach for DNA extraction from FFPE tissues

**DOI:** 10.1101/2024.07.08.602387

**Authors:** Christoph Geisenberger, Edgar Chimal, Philipp Jurmeister, Frederick Klauschen

**Affiliations:** Institute of Pathology, Ludwig-Maximilians-Universität München, Munich, Germany; German Cancer Consortium (DKTK), partner site Munich, a partnership between DKFZ and University Hospital Munich, Germany; BIFOLD – Berlin Institute for the Foundations of Learning and Data, Berlin, Germany

**Keywords:** FFPE, DNA extraction, DNA methylation, molecular pathology

## Abstract

Genomic profiling of cancer plays an increasingly vital role for diagnosis and therapy planning. In addition, research of novel diagnostic applications such as DNA methylation profiling requires large training and validation cohorts. Currently, most diagnostic cases processed in pathology departments are stored as formalin-fixed and paraffin embedded tissue blocks (FFPE). Consequently, there is a growing demand for high-throughput extraction of nucleic acids from FFPE tissue samples. While proprietary kits are available, they are expensive and offer little flexibility. Here, we present ht-HiTE, a high-throughput implementation of a recently published and highly efficient DNA extraction protocol. This approach enables manual and automated processing of 96-well plates with a liquid handler, offers two options for purification and utilizes off-the-shelf reagents. Finally, we show that methylation profiles obtained from DNA processed with ht-HiTE are of equivalent quality as compared to a manual, kit-based approach.

## Introduction

The introduction of tissue preservation with formalin in the late 19th century allowed researchers to conserve a variety of samples for extended periods without significant degradation ^1^. Formalin fixation followed by paraffin embedding (FFPE) is still the gold standard for the storage of tissue specimens in diagnostic pathology. FFPE tissue blocks can be kept at room temperature and processing them as tissue sections for (immuno)histochemistry is straightforward. While biomolecules such as nucleic acids and proteins are maintained, formalin fixation and extended storage lead to crosslinking, fragmentation and other types of damage (reviewed in Steiert et al.^2^). Sequencing RNA from FFPE tissue is especially challenging ^3^. However, technological progress in sequencing library generation has resulted in much improved sensitivity. It is now possible to generate whole genome and whole exome sequencing libraries from a few nanograms of DNA and multiple research groups have achieved single-cell RNA sequencing (scRNA-seq) in FFPE tissues ^4,5^. At the same time, the demand for high-throughput analysis of nucleic acids in FFPE tissues has grown rapidly. This is mostly due to the increased use of next generation sequencing in cancer. Profiling of single-nucleotide variants (SNVs), gene fusions and somatic copy number aberrations (SCNA) is required more and more for the diagnostic workup and therapy planning of tumor patients. Furthermore, new diagnostic applications in molecular pathology, such as DNA methylation classification, are showing promise ^6–10^. These methods use machine learning and artificial intelligence and therefore require large training and validation cohorts. To scale up mutation and DNA methylation profiling, high-throughput DNA extraction is necessary, ideally from FFPE tissues where most tumor samples are stored. Commercial applications are offered by a number of companies such as Maxwell (HT DNA FFPE) and Covaris (truXTRAC® FFPE SMART). However, they use proprietary reagents and are expensive. This precludes researchers from making informed changes to the protocol which may be necessary for specific applications. In addition, early prototyping and testing can benefit from more cost-efficient protocols. Recently, Oba and colleagues provided an interesting new approach for DNA extraction from FFPE tissues ^11^. Termed HiTE (Highly concentrated Tris-mediated DNA extraction), their approach utilizes high concentrations of the formalin scavenger Tris (tris[hydroxymethyl]aminomethane). This resulted in higher yields and DNA quality, presumably due to more efficient de-crosslinking. This was also reflected in improved data quality in NGS experiments. This manuscript extends HiTE to a high-throughput format (ht-HiTE) that can be performed manually or using liquid handlers. Also, we demonstrate that DNA methylation profiling of samples processed with ht-HiTE produces robust, high-quality data comparable to a kit-based manual workflow. Lastly, we offer the research community a detailed online version of the protocol.

## Results

### Overview of the protocol and manual testing

The high-throughput implementation presented here is based on HiTE, a method published by Oba and colleagues^11^. The protocol includes four basic steps: (i) deparaffinization, (ii) tissue lysis, (iii) reversal of interstrand crosslinks and (iv) DNA purification. Single Eppendorf tubes or 96-well deep well plates are pre-loaded with 500 µl mineral oil per reaction chamber. Then, tissue scrolls or fragments scratched from glass slides are placed in each tube or well. Paraffin is removed by immersion in mineral oil and melting, followed by Proteinase K digestion. De-crosslinking is performed at 80°C overnight in the presence of 800 µM Tris. Finally, DNA can be purified using silica columns or paramagnetic SPRI beads. First, we assessed the performance of manual HiTE in our hands. Whole tissue scrolls (10 µM) from an FFPE sample (1 year old) were processed manually in replicates of four per condition. As a reference, we extracted DNA with the Maxwell® CSC DNA FFPE Kit according to the manufacturer’s instructions (from here on referred to as *gold standard*). HiTE samples were processed as outlined above and purified either with Ampure XP SPRI beads or Quiagen silica columns. Also, we assessed two different incubation times: 24 hours as suggested by Oba et al. and 1 hour. After purification, yields were quantified with a dye-based method (Qubit^™^) or spectrometrically (Nanodrop^™^). Longer incubation yielded significantly more DNA as measured by Qubit (Fig. 1a; p = 2 × 10^−5^, *Student’s t-test*). At 24 hours, the average yield was 15.8 ng/ul with similar results for HiTE and the gold standard (Fig 1a; p > 0.65, *ANOVA*). Purity as measured by the A260/A280 ratio was 1.8 for column-based and 1.7 for bead-based purification (Fig. 1b). For the gold standard, purity showed slightly divergent results depending on the incubation time with a ratio of 1.87 after 1 hour and 1.76 after 24 hours. Plotting the concentrations measured by Nanodrop and Qubit against each other revealed a strong linear relationship (Fig. 1c). However, as is well known, Nanodrop measurements overestimate DNA content with elevated overestimation for the gold standard after 1 hour of incubation. Excluding the outliers (n = 4) revealed a correlation of 0.95 (Pearson’s *r*) between measurements (r = 0.63 when including all samples). Nanodrop overestimation was roughly 2-fold (Fig. 1d) with a concentration-dependent effect which reached a plateau for concentrations of >10 ng/µl (Qubit) or >20 ng/ul (Nanodrop) (Fig. 1e). Taken together, these findings validate HiTE as an useful approach for high-quality DNA extraction with off-the shelf reagents.

**Figure 1.**
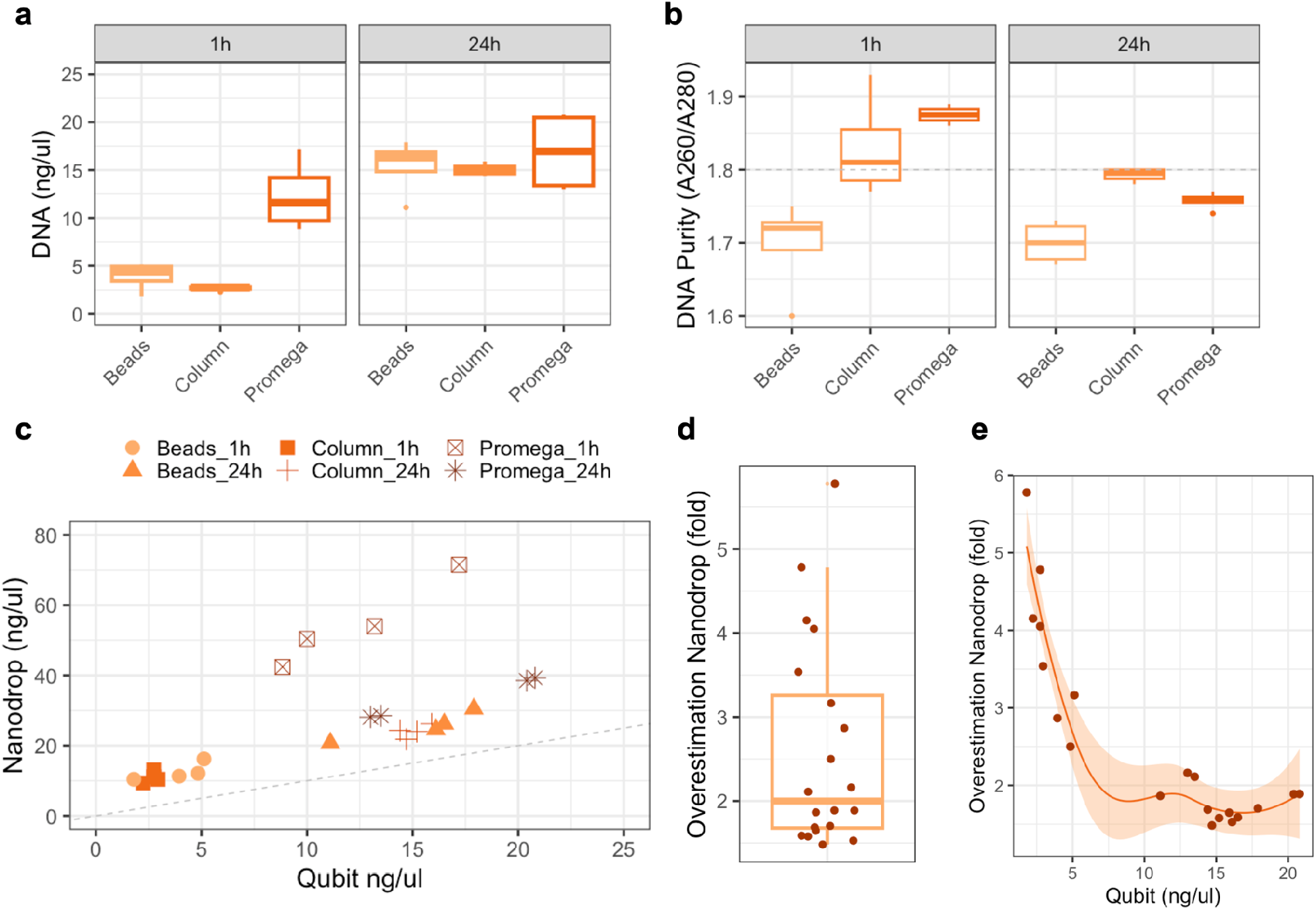
Manual validation of highly concentrated Tris-mediated DNA extraction (HiTE) **a**. shows DNA concentrations obtained after 1 hour (left panel) and 24 hours (right panel) of incubation. Samples processed with off-the-shelf reagents and purified with beads or columns yielded roughly equal amounts compared to the “gold standard” (Promega kit). **b**. shows the ratio of absorption at 260 and 280 nm based on spectrometry reading. A ratio of 1.8 (dashed grey line) is considered pure for DNA samples. **c**. DNA concentrations for all samples were measured with a spectrometry-based (Nanodrop) and dye-based (Qubit) approach. Except for the gold standard after 1 hour of incubation, there is a strong linear relationship between both measurements with overestimation of yield by spectrometry. This overestimation is roughly two-fold (**e**) and depends on the amount of DNA (**f**). In general, deviation is larger for smaller DNA concentrations.

### Plate-based processing

Next, we established an automated version of the protocol on a liquid handler (Beckman Coulter Biomek i5). A total of 92 samples and 4 empty controls were processed in duplicate in two 96-well plates. Processing was identical to the manual samples and DNA extraction was performed with either SPRI beads or columns for one plate each (Fig. 2a). Yield as measured by DNA concentration showed much larger variation than observed for manual processing, owing to the larger differences in tissue size between the processed blocks. Mean and median Qubit readings were 86.6 and 72.0 ng/µl across all non-control samples (Fig. 2b, Fig S1a). Stratifying for the manner of purification revealed small but significant differences between approaches (Fig. 2c). Bead-based purification yielded on average 97.0 ng/µl compared to 68.6 ng/µl for columns as measured by Qubit (p = 0.023, *Welch’s two-sided t-test*). Nanodrop based measurements were 234.8 ng/µl and 157.6 ng/µl, respectively (p = 0.009, *Welch’s two-sided t-test*). DNA purity as measured by the A260/A280 ratio was 1.84 for bead-based and 1.9 for column-based purification (Fig. 2d). Controls had significantly lower readings with zero measurements for the majority of empty wells (p = 6.7 × 10^−28^, *Welch’s two-sided t-test*, Fig. S1b). These results highlight that HiTE can successfully be implemented in an automated setting.

**Figure 2.**
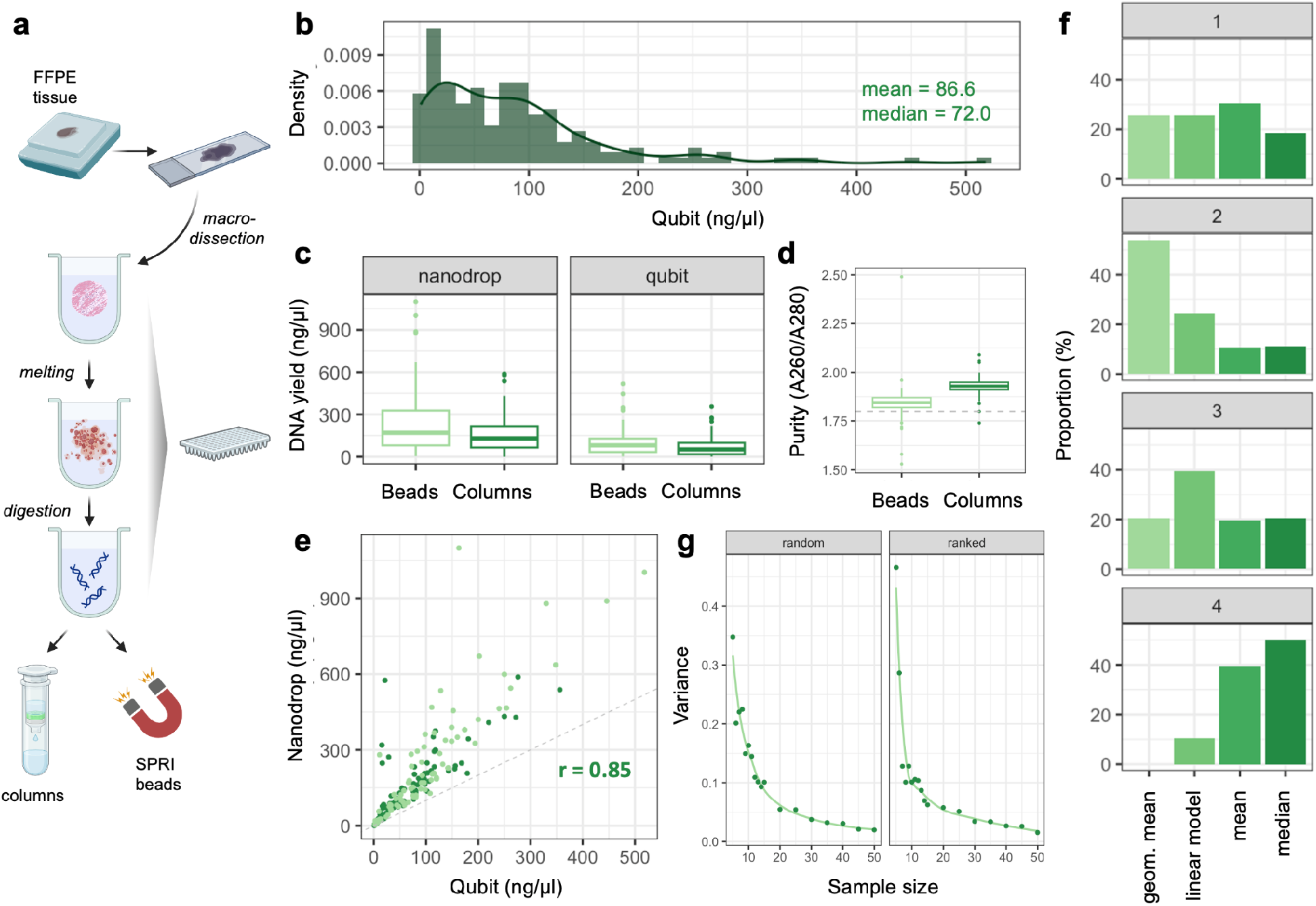
Plate-based processing and correction of spectrometry-based readings. **a**. Overview of sample processing **b**. Distribution of DNA content for two 96-well plate measured with Qubit **c**. Difference in concentrations between the plates loaded with replicates. Bead-based purification yielded slightly but significantly higher DNA amounts. **d**. UV absorption ratio (260 nm / 280) nm absorption ratio was closer to 1.8 for samples purified with SPRI beads. **e**. Scatterplot for dye- and spectrometry-based measurements. There is high correlation between measurement approaches (Pearson’s r = 0.85) with roughly two-fold overestimation by Nanodrop **f**. Provides a ranking for four different approaches for correcting Nanodrop overestimation, based on ranking mean-squared error (MSE) across 200 random splits of the data. Panels indicate how many times each approach scored first (*top panel*), second, third or last (*bottom panel*). Overall, correction by geometric mean performed best. **g**. Provides an estimate of the variance observed (y-axis) when estimating the geometric mean for a limited number of samples (x-axis). Samples were either picked randomly (*left panel*) or evenly spaced according to their DNA concentrations (*right panel*). The elbow indicates that approximately 20 samples are sufficient for a stable estimation of the geometric mean

### Correction of spectrometry-based DNA measurements

Dye-based measurements are more specific and sensitive than UV-based approaches, but carry a higher price point. Comparing both measurements (Fig. 2e), we noted good agreement with a correlation of 0.85 (*Pearson’s r*). However, spectrometry-based measurements again exhibited a roughly 2-fold overestimation with larger deviations for lower DNA content (Fig. S1c). We therefore tested how well a simple correction factor estimated from a subset of wells can ameliorate overestimation. Randomly splitting the data into a training and test cohort 200 times, we fitted a linear model and also calculated the median, mean or geometric mean for the overestimation observed in the training cohort. Then, each approach was used to correct the concentrations of the test samples and the overall performance was ranked by the mean squared error (MSE). Next, performance was compared by calculating how many times each correction factor calculation scored first, second, third of last (Fig. 2f). Overall, a simple correction factor based on the geometric mean performed best and was selected for subsequent analyses. Next, we assessed how many samples are required to obtain a reliable estimate of the correction factor (i.e., geometric mean). A subset of samples was selected and used to calculate Nanodrop overestimation. The variance of the estimate decreased with sample size and showed an elbow for sample sizes of 15 to 20 (Fig. 2g, Fig. S2). This was observed both for samples selected randomly (left panel) or spaced evenly according to Nanodrop measurement (right panel) with lower variances observed for the latter. These results (i) indicate that picking samples based on their concentration performs better and (ii) that samples size of 20 are sufficient to obtain reliable estimates of a factor to correct spectrometry-based reading. We applied the strategy outlined above for the two plates processed in this study. Inferred concentrations deviated less than two-fold in either direction (i.e., 0.5x or 2x) for 94.0% and 89.3% of cases in bead- and column-based samples, respectively.

### Methylation profiling of samples processed in 96-well format

To assess the quality of DNA generated with ht-HiTE, methylation data were generated in triplicate for three FFPE blocks (9 to 12 years old). For each case, technical replicates were generated with DNA from (i) manual processing, (ii) ht-HiTE with column purification and (iii) ht-HiTE with bead purification. First, basic quality measures were assessed. Detection rate, i.e. the proportion of probes on the array which yielded usable data, was >99.9% for all samples (Fig. 3a) with a trend towards less failed probes for column purification. Relative methylation as measured by the beta value (methylated signal divided by sum of methylated and unmethylated signal, M / [U + M]) is bounded between 0 and 1 and typically shows a bimodal distribution. The same was observed when plotting the distribution of beta values across samples (Fig. 3b) without major differences between processing modalities. Selecting the 5,000 most variable probes across all samples, clustering of beta values revealed the highest similarities between DNAs from the same tissue irrespective of purification (Fig. 3c). Correlation of beta values was very high for technical replicates (Fig. 3d, 0.96 to 0.98, Pearson’s *r*) and higher than between-sample correlations (0.89 to 0.94, Pearson’s *r*). Unsurprisingly, clustering samples based on their pairwise correlations reproduced the high similarity for technical replicates (Fig. 3e and Fig. S3). Finally, genome-wide copy-number plots generated with the software *conumee* ^12^ showed highly reproducible profiles between DNAs extracted by different means (Fig. 3f, Fig. S4 and Fig. S5).

**Figure 3.**
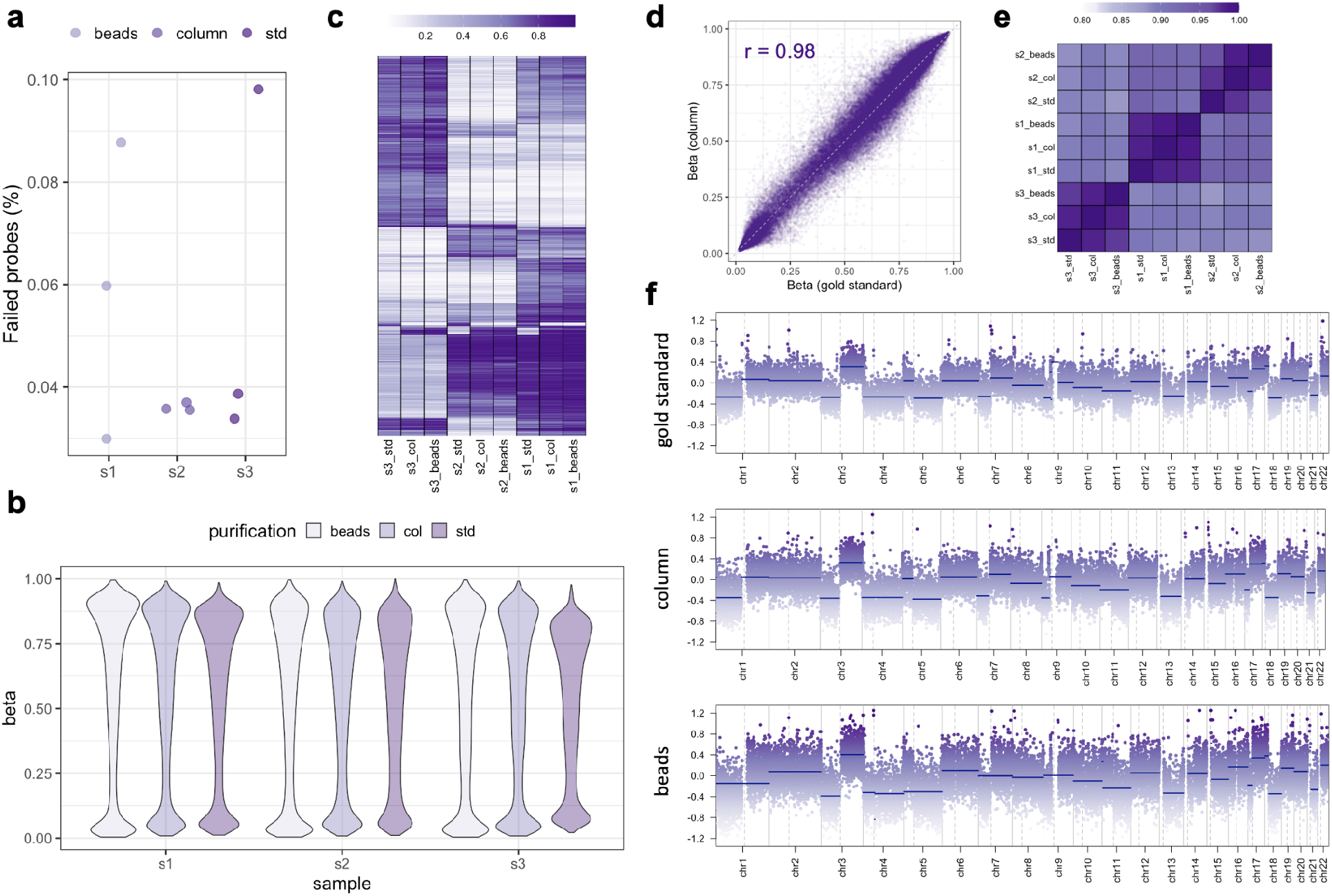
Comparison of methylation data for manual and plate-based processing. **a**. shows the amount of failed probes (detection p-value > 0.05). While ht-HiTE, column-cleaned reactions had the lowest probe failure, all samples exhibited detection rates of > 99.9%. **b**. Provides an overview of the distribution of beta values as violin plots for all samples. As expected, values are centered around 1 (full methylation) and 0 (methylation absent) without major differences between purification approaches. **c**. shows a heatmap for the 5,000 most variable probes (rows) on the array with darker colors representing higher methylation. Samples cluster primarily by patient (columns). **d**. Example scatterplot of beta values for sample 1, comparing manual gold standard (x-axis) and high-throughput column cleanup (y-axis). Reproducibility is high with a correlation (Pearson’s r) of 0.98. Different purification approaches the gold standard (x-axis) vs. a high-throughput, column-cleaned sample (y-axis). **e**. Heatmap for all pair-wise correlations. X- and y-axis are clustered with hierarchical clustering (dendrogram not shown). Within-sample correlations are higher than between-sample correlations. **f**. shows copy-number plots extracted from methylation array data. Here, the observed alterations can reliably be detected for manual / gold standard (top), plate-based / column (middle) and plate-based / beads (bottom) purification

## Discussion

In this manuscript, we presented and thoroughly tested ht-HiTE, the high-throughput implementation of HiTE, a technique for DNA extraction published by Oba and colleagues ^11^. Manual testing validated that (ht-)HiTE yields significant amounts of DNA with good purity, comparable to the Maxwell® CSC DNA FFPE Kit. Extended incubation times (24 hours) significantly increased DNA yield compared to shorter times (1 hour). DNA purity, measured by the A260/A280 ratio, was similar between ht-HiTE and the gold standard, confirming the method’s reliability. In addition, we provided evidence that purification using SPRI beads is effective and reliable. Automating the ht-HiTE protocol on the Beckman Coulter Biomek i5 platform demonstrated its scalability for high-throughput applications. Although DNA yield varied more in automated processing due to differences in tissue size, bead-based purification consistently yielded higher DNA concentrations than column-based purification. The samples processed in this study encompass different subtypes of sarcoma, including calcified and adipose tissue, which underscores the broad applicability in terms of tissue types.

A significant challenge encountered was the overestimation of DNA content by UV spectrometry (Nanodrop) compared to dye-based methods (Qubit). Nanodrop measurements overestimated DNA content by approximately two-fold, especially for lower concentrations. Applying a correction factor based on the geometric mean of overestimation improved the accuracy of DNA quantification. A sample size of around 20 was sufficient to obtain reliable estimates. Measuring only a subset of samples allows researchers to further lower the price point. Omitting the RNAse digest and dilution of SPRI beads with lab-made bead binding buffer allows DNA extraction for as little as 2 euros per sample based on list prices in Germany as of May 2024 (Table 1).

**Table 1.** Cost comparison of plate-based purification. Using the prices available in Germany as of May 2024, processing costs per sample were calculated for column-based (left) and bead-based (right) purification. Lowest prices can be achieved for bead purification without RNAse treatment, dilution of bead-binding buffer and obtaining dye-based measurements for only a subset of samples.

| Reagent | Price per sample<br>(columns) | Price per sample<br>(beads) |
| --- | --- | --- |
| Glass slides | € 0.64 | € 0.64 |
| Water | € 0.03 | € 0.03 |
| 96-well plates | € 0.10 | € 0.10 |
| Mineral oil | € 0.10 | € 0.10 |
| Tris-HCl | € 0.16 | € 0.16 |
| SDS 10% | € 0.01 | € 0.01 |
| Proteinase K | € 0.24 | € 0.24 |
| RNAse A | € 0.32 | € 0.32 |
| Spin columns (96w) | € 3.63 |  |
| SPRI beads |  | € 1.01 |
| Qubit dsDNA Kit | € 0.80 | € 0.80 |
| Total: | € 6.02 | € 3.41 |
| Without RNAse: | € 5.70 | € 3.09 |
| No RNAse, diluted<br>beads, Nanodrop<br>correction | € 5.06 | € 1.69 |

Methylation profiling further validated the quality of DNA extracted using ht-HiTE. Methylation data from manually and ht-HiTE processed DNA showed high detection rates and consistent beta value distributions. Clustering of the most variable probes and correlation analysis confirmed the high similarity between DNAs extracted by different methods, indicating that the ht-HiTE protocol maintains DNA integrity. Of note, gold standard samples were assayed on a different array platform (EPIC v1) as the ht-HiTE samples (EPIC v2). Nevertheless, correlations between technical replicates are as high as those reported in the characterization of the novel EPIC v2 platform ^13^.

These findings have significant implications for molecular pathology, particularly in high-throughput DNA extraction and analysis. The ht-HiTE protocol offers a cost-effective, scalable solution for extracting DNA from FFPE tissues, facilitating next-generation sequencing and other high-throughput techniques in cancer diagnostics and research. In summary, the ht-HiTE protocol for DNA extraction from FFPE tissues is robust and reliable for both manual and automated settings. Its ability to produce high-quality DNA suitable for downstream applications makes ht-HiTE a valuable tool for molecular pathology. Future studies should aim to optimize the protocol further and explore its applicability to various tissue types and clinical samples.

## Materials and Methods

### Ethics Statement

The research project has been approved by the ethics committee of LMU University Munich. All analyses were retrospective and conducted with leftover material of diagnostic cases.

### Code and Data Availability

Raw DNA microarray data generated in this study are available at figshare (http://dx.doi.org/10.6084/m9.figshare.26198033). Code to reproduce results and generate plots is available on github (https://github.com/cgeisenberger/ht-hite).

### Experimental Methods Availability

A detailed step-by-step version of the protocol has been published at protocols.io and is accessible at dx.doi.org/10.17504/protocols.io.6qpvr3jr3vmk/v1.

### Study Design

The main research objective of this study was to establish a high-throughput implementation of a FFPE DNA extraction protocol using high concentrations of Tris (ht-HiTE). The workflow was tested manually and compared to the kit-based gold standard used in the lab. Next, an automated version was set up to process replicates deposited in two 96-well plates. Finally, DNA microarray profiling of DNAs extracted with the automated workflow were compared to previously generated data to assess the quality of methylation data attainable with ht-HiTE. We obtained

### Patients and Samples

Samples were selected from the archives of the Institute of Pathology at LMU Munich. Manual testing was performed from replicates of a colorectal cancer sample processed in 2023. Plate-based processing was performed for different subtypes of sarcoma with tissue block ages ranging from 1 to 7 years old (n = 89), lung cancer samples between 9 and 12 years old (n = 3) and empty controls (n = 4). Methylation profiles were generated for lung cancer samples processed with ht-HiTE and cleaned with beads or columns (3 samples with 2 replicates, n = 6 total). These data were compared to previous array experiments of DNA extracted with the gold standard.

### DNA extraction: Gold standard

Tissue sections (2 µm) were stained with hematoxylin and eosin (H&E) to select regions with high tumor cell content. Macroscopic dissection of tumor areas was performed with sterile scalpels and tissues were extracted using the Maxwell RSC FFPE Plus DNA Purification Kit (Promega) according to the manufacturer’s instructions.

### DNA extraction: Automated HiTE

Tissue sections (2 µm) and unstained sections (10 µm) were placed on glass slides (TOMO, TOM-14). After staining 2 µm sections with H&E, areas of interest were macroscopically dissected with a scalpel blade (for example Ruck, 2009010) and placed in 1 ml DNA lo-bind, deep-well 96-well plates (Eppendorf, 0030503244) preloaded with 500 µl of mineral oil (Sigma Aldrich, M5904-500ML). Plates were incubated for 15 min at 56°C in a thermocycler to melt paraffin. Next, 100 µl of the following mix were added to each sample: 80 µl Tris-HCl pH 8.0 (Merck Millipore, 648314-100ML), 10 µl SDS 10% (Sigma Aldrich, 71736-100ML), 5 µl Proteinase K (NEB, NEB, P8107S), 5 µl H_2_O (Promega, P1199). Samples were incubated 1 hour at 56°C followed by 16 to 24 hours at 80°C and a holding step at 4°C. For bead-based purification, samples were transferred to a fresh, shallow 96-well plate (Eppendorf, 0030503104), mixed with 80 µl of Ampure XP beads (Beckman Coulter, A63882) and incubated for 10 minutes at room temperature. Next, samples were placed on a magnetic rack to pellet beads. Then, supernatant was aspirated and 200 µl of 70% ethanol were added to each well. After 30 s to 1 min, ethanol was removed. After repeating the wash step once, the beads were dried at room temperature for 10 minutes. Next, the plate was removed from the magnet and 60 µl of nuclease free water was added to each sample. Liquid and beads were mixed by pipetting and incubated for 10 minutes at room temperature off the magnet, Finally, the plate was placed back on the magnetic rack and the purified DNA sample was transferred to a fresh DNA lo-bind plate. For column-based purification, sample volume was adjusted to 200 µl with nuclease-free water. Next, purification was performed using the DNAeasy Blood and Tissue spin column kit (Qiagen, 69581) according to the manufacturer’s instructions. Briefly, 400 µl of Buffer AL was added to each sample and the full volume was transferred to a spin plate. After centrifugation and two cleaning steps with Buffer AW1 and AW2, samples were eluted in 60 µl of buffer AE. The steps outlined above were automated on a Biomek i5 liquid handling station (Beckman Coulter). Of note a number of samples (n = 8) evaporated during DNA extraction due to imperfect sealing and were excluded from the analysis.

### DNA quantification

Nucleic acids were quantified using the dye-based Qubit^™^ HS DNA Assay (Thermo Fisher) or spectrometrically with the NanoDrop^™^ One platform (Thermo Fisher).

### Microarray data processing and statistical analysis

Microarray methylation data were processed using the software package *minfi* ^*14*^ and normalized using single-sample normal-exponential out-of-band (Noob) normalization ^15^. Downstream analysis was performed using the software package *tidyverse*. Copy-number plots were generated using *conumee2*.*0*^*12*^. Of note, manually extracted DNA samples were assayed on a different microarray platform (EPIC v1) as ht-HiTE samples (EPIC v2). Data were combined by selecting probes available on both platforms (n = 718,960). Normal control samples for copy number plots (n = 5 for each array type) were downloaded from the NCBI GEO database available under accession GSE235717 (EPIC v1) and GSE246337 (EPIC v2).

