## Supplemental Figures for "A cost-effective and scalable approach for DNA extraction from FFPE tissues"

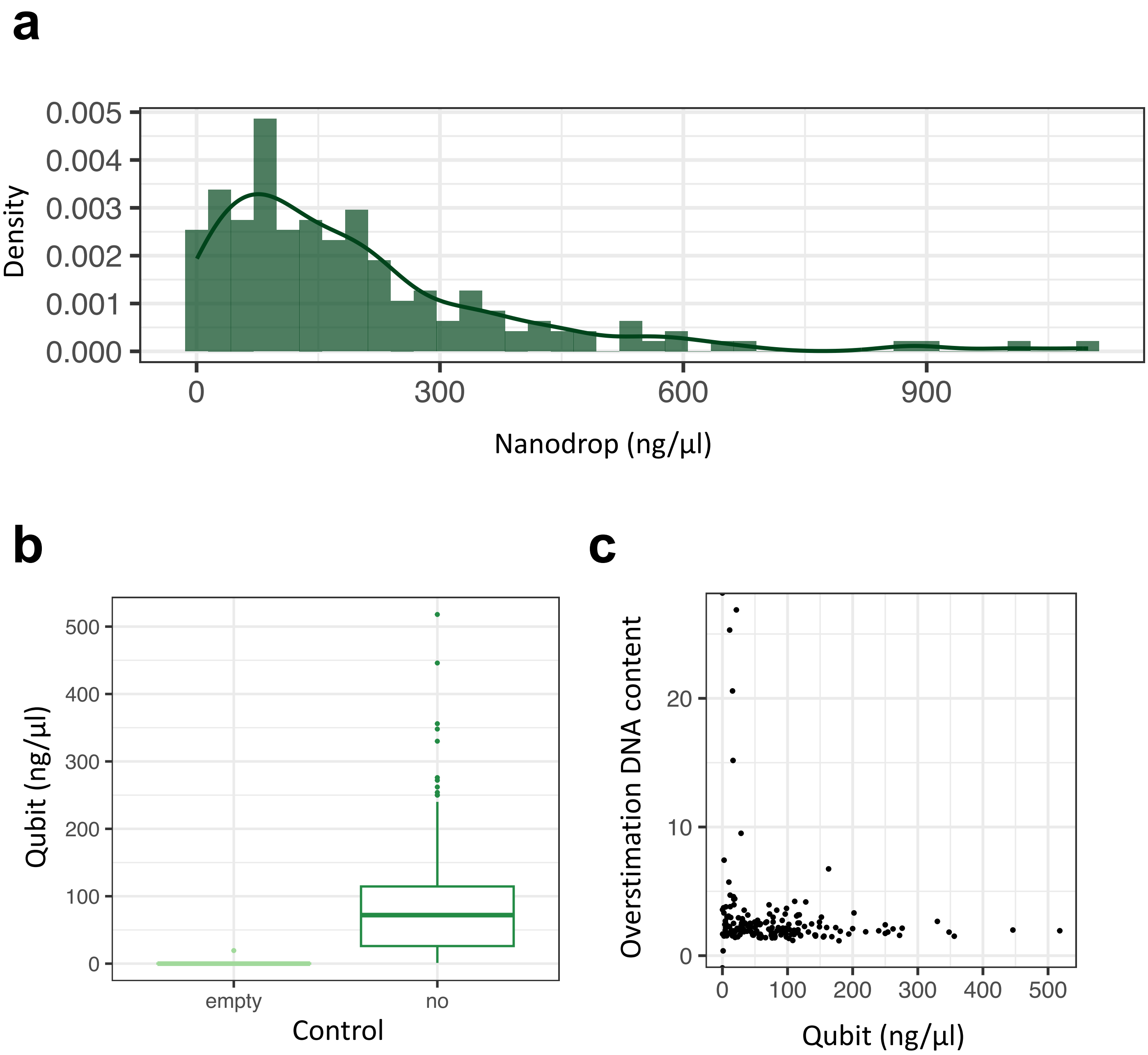


**Figure S1. Implementation of plate-based HiTE**

**a.** shows a density plot for concentrations as measured by Nanodrop for two plates processed with ht-HiTE. **b.** compares Qubit readings for empty controls and regular samples for two ht-HiTE plates. **c.** Shows the overestimation of DNA yield as measured by Nanodrop (y-axis) in relation to the actual amount of DNA as measured by Qubit (x-axis).

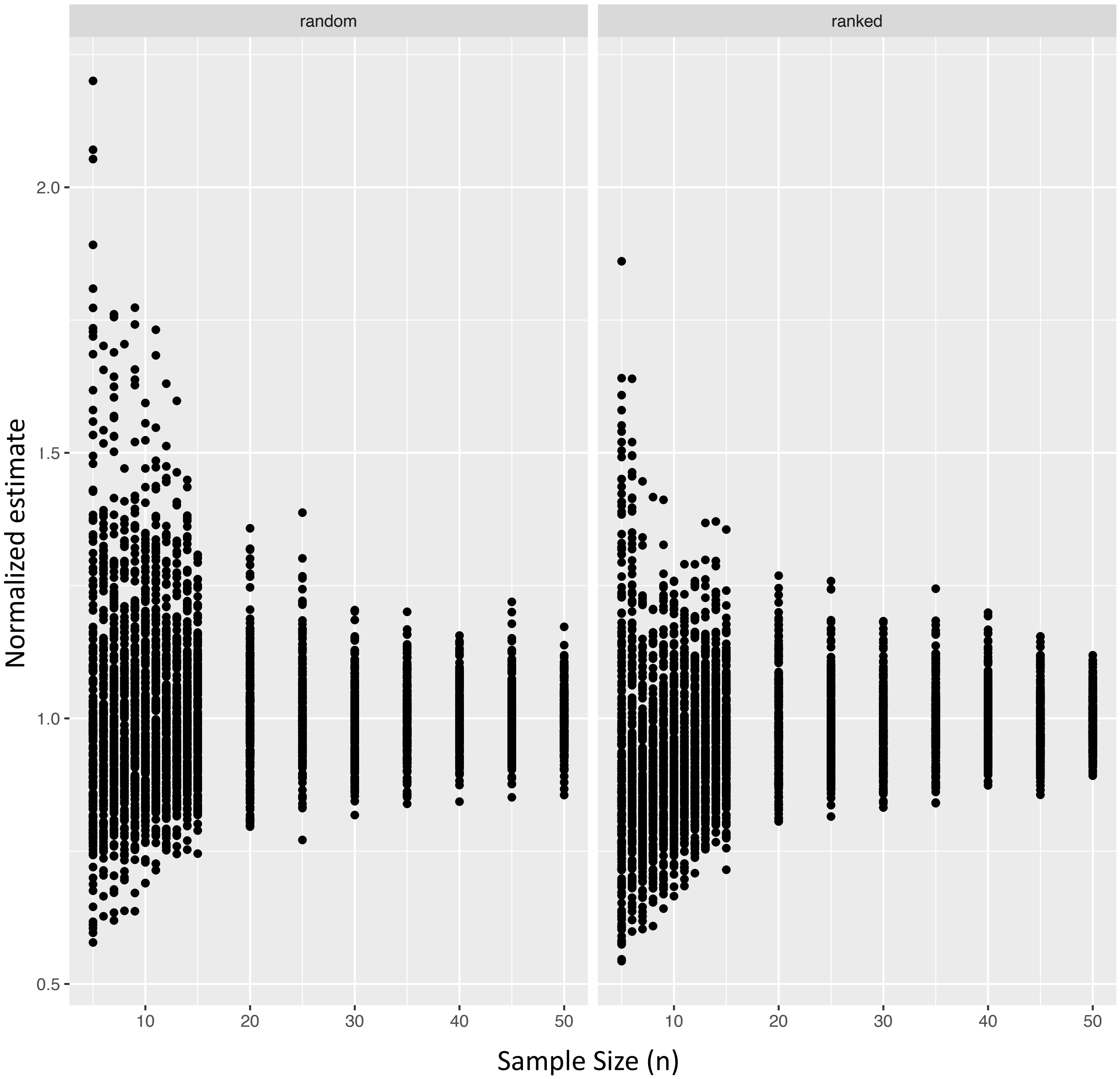


**Figure S2. Estimates of geometric mean for different sample sizes**
Increasing sample sizes (x-axis) were used to estimate the geometric mean of DNA content overstimation (ratio Nanodrop/Qubit reading, y-axis). For each sample size, 200 bootstraps of the data (80%) were performed and the overestimation was normalized by dividing through the actual value per bootstrap. Samples were either sampled randomly (left panel) or ranked according to the Nanodrop reading (right).

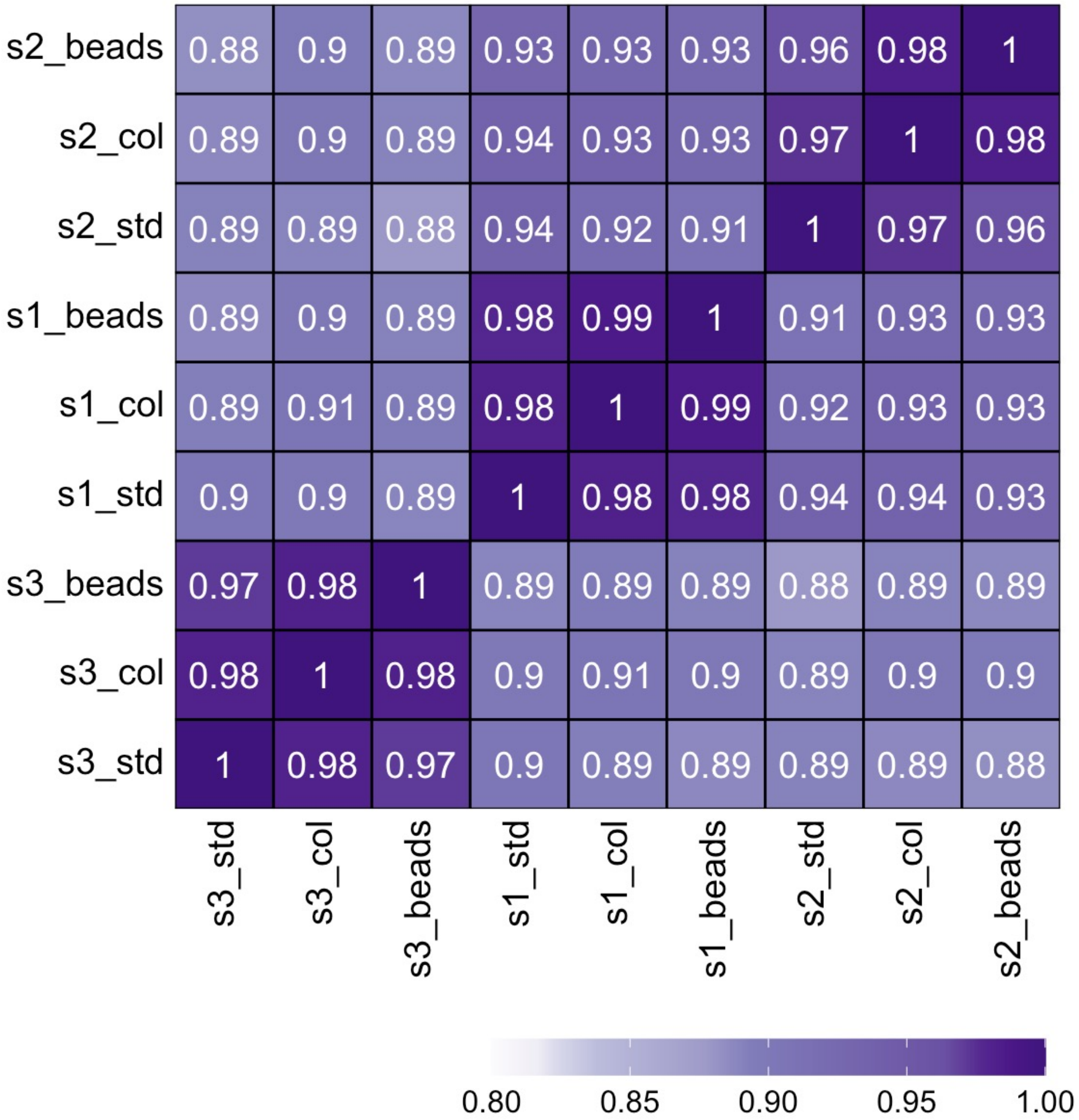


**Figure S3. Heatmap of pairwise beta value correlations**

Similar to Fig. 3e, pairwise correlations for all samples are shown with the actual value (Pearson's *r*) added to the plot.

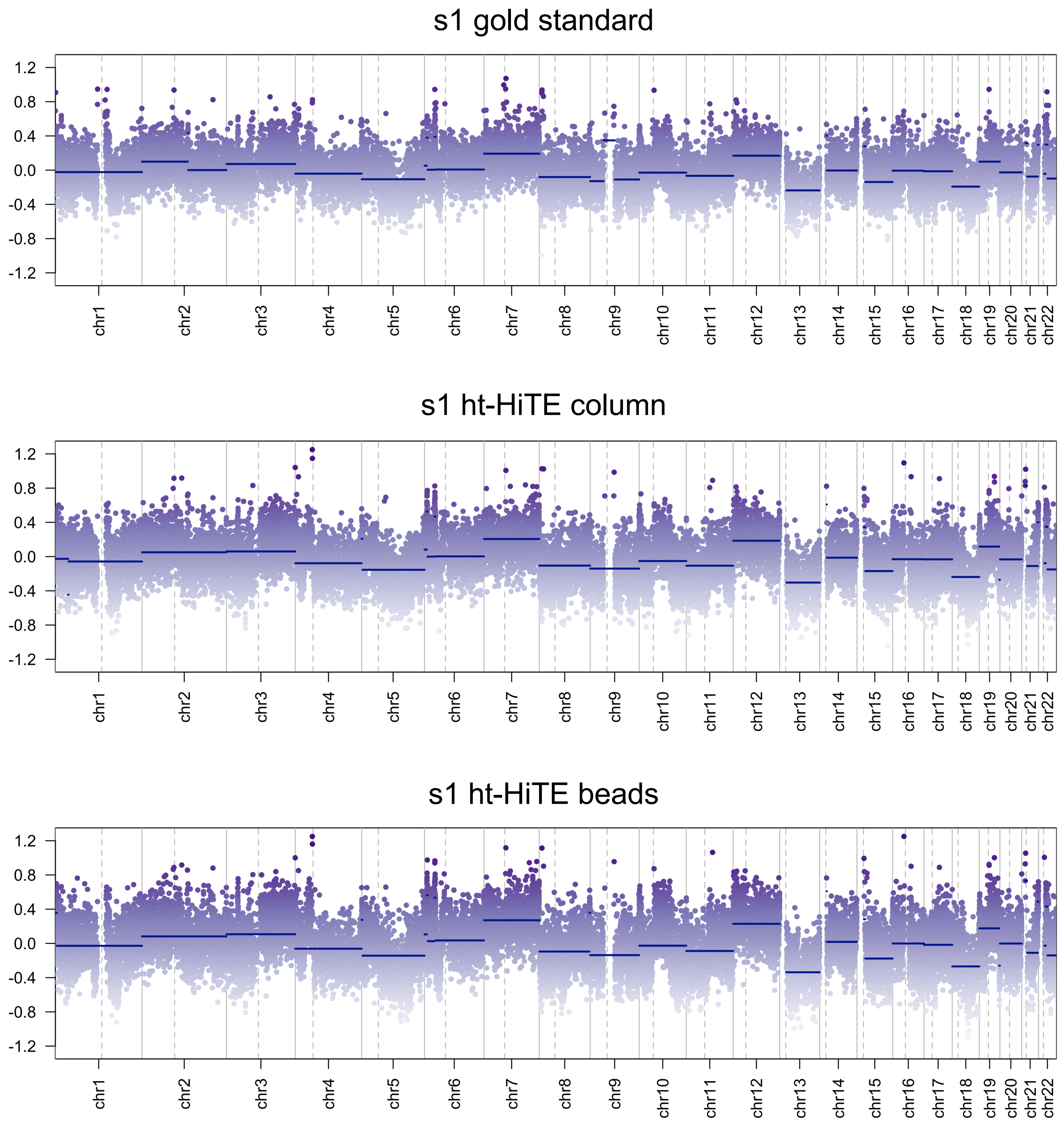


**Figure S4. Copy number plots for sample 1**

Copy number plots for the manual DNA preparation (top), ht-HiTE with column purification (middle) and ht-HiTE with bead purification (bottom).

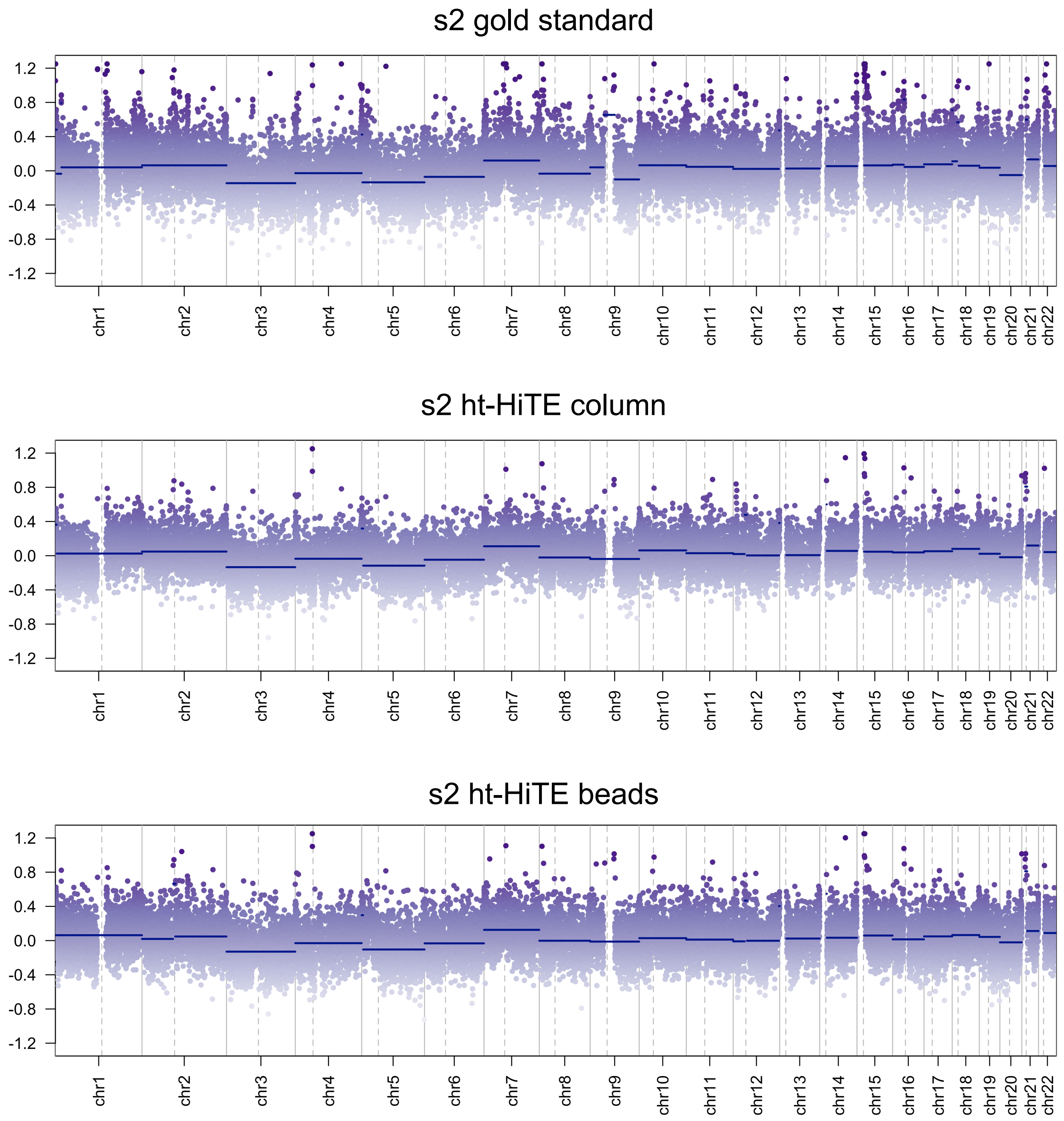


**Figure S5. Copy number plots for sample 2**

Copy number plots for the manual DNA preparation (top), ht-HiTE with column purification (middle) and ht-HiTE with bead purification (bottom).
